# Diet, psychosocial stress, and Alzheimer’s disease-related neuroanatomy in female nonhuman primates

**DOI:** 10.1101/2020.09.25.313593

**Authors:** Brett M. Frye, Suzanne Craft, Thomas C. Register, Rachel N. Andrews, Susan E. Appt, Mara Z. Vitolins, Beth Uberseder, Marnie G. Silverstein-Metzler, Haiying Chen, Christopher T. Whitlow, Jeongchul Kim, Richard A. Barcus, Samuel N. Lockhart, Siobhan Hoscheidt, Brandon M. Say, Sarah E. Corbitt, Carol A. Shively

**Author notes:** Corresponding Author: Carol A. Shively, PhD, Department of Pathology/Comparative Medicine, Wake Forest School of Medicine, Medical Center Blvd, Winston-Salem, NC 27157-1040.

## Abstract

**INTRODUCTION:** Associations between diet, psychosocial stress, and neurodegenerative disease, including Alzheimer’s disease (AD), have been reported, but causal relationships are difficult to determine in human studies.

**METHODS:** We used structural magnetic resonance imaging in a well-validated nonhuman primate model of AD-like neuropathology to examine the longitudinal effects of diet (Mediterranean versus Western) and social subordination stress on brain anatomy, including global volumes, cortical thicknesses and volumes, and twenty individual regions of interest (ROIs).

**RESULTS:** Western diet resulted in greater cortical thicknesses, total brain volumes and gray matter, and diminished cerebrospinal fluid and white matter volumes. Socially stressed subordinates had smaller whole brain volumes but larger ROIs relevant to AD than dominants.

**DISCUSSION:** The observation of increased size of AD-related brain areas is consistent with similar reports of mid-life volume increases predicting increased AD risk later in life. While the biological mechanisms underlying the findings require future investigation, these observations suggest that Western diet and psychosocial stress instigate pathologic changes that increase risk of AD-associated neuropathologies, whereas Mediterranean diet may protect the brain.

**RESEARCH IN CONTEXT:**

1. Systematic review: The authors reviewed the literature with PubMed and Google Scholar and found a number of publications which are cited that suggest that AD pathogenesis begins well before the onset of symptoms.
2. Interpretation: Our findings support the hypothesis that Western diet and psychosocial stress may instigate neuroinflammatory responses that increase risk of later developing AD-like neuropathologies, whereas the structural stasis in the Mediterranean diet group may represent a resilient phenotype.
3. Future directions: The manuscript serves as a critical first step in describing risk and resilient phenotypes during middle age in a nonhuman primate model of AD-like neuropathology. This report lays the groundwork for ongoing efforts to determine whether neuroinflammatory profiles differed across diet and stress groups. Future studies should aim to understand the temporal emergence of functional disparities associated with the changes in brain structure observed here.

**HIGHLIGHTS:**

- Global brain volumes changed in response to Western, but not Mediterranean, diet.
- Western diet increased cortical thickness in multiple regions relevant to AD.
- Mediterranean diet did not alter cortical thicknesses relevant to AD.
- Brain regions associated with AD risk differed between low and high stress monkeys.
- Psychosocial stress may modulate the effects of diet on the brain.

## INTRODUCTION

Diet composition may be an important factor mediating risk of neurodegenerative disease [1]. Routine consumption of a Western diet – high in saturated animal fats, simple sugars, and sodium – increases risk of obesity, diabetes, and autoimmune and cardiovascular diseases [2-4], all of which increase AD risk [5]. In observational studies, Western diet consumption is associated with neuroanatomical changes, e.g., reductions in hippocampal volume, that are associated with diminished cognitive performance [6]. In contrast, Mediterranean diet consumption is associated with healthy aging outcomes [7]. Adherence to Mediterranean-type diets reduces circulating biomarkers of ischemic heart disease risk [17] and is associated with decreased risk of cardiovascular disease [4], cognitive impairment [8], and AD [9]. While plausible biological mechanisms of these effects have been identified [10,11], most of these data derive from observational studies in which dietary consumption is self-reported. Likewise, those who report adhering to Mediterranean versus Western diet patterns are different in other ways that may affect cognitive health (e.g., exercise, education, smoking behavior), making the process determining causal factors difficult.

Psychosocial stress is associated with increased risk of AD and other dementias [12]. Socioeconomic disparities are one of the clearest drivers of chronic psychosocial stress, and the gradient of health disparities resulting from stress may become steeper with age [13]. Low socioeconomic status (SES) increases risk of dementia, but the factors underlying this disparity are not well understood [14]. A number of pathways which are affected by stress that contribute to neurodegeneration and cognitive decline have been identified [15]. However, those high and low in SES differ in a number of ways that may influence health (e.g. physical activity, diet, education, income, neighborhood, reserve) complicating the process of uncovering mechanisms. Likewise, the effects of psychosocial stress may be temporally distant from the stressor and often are based on retrospective self-report [12,16].

Understanding how dietary patterns and stress profiles modify AD risk is key to identifying therapeutic targets. However, randomized clinical trials of long-term effects of diet are difficult and expensive, and randomized clinical trials of psychosocial stress effects on health are unethical. While transgenic rodent models have provided invaluable mechanistic insights into early-onset AD, they have been less helpful in understanding late-onset sporadic AD which accounts for about 95% of cases [17].

Macaques share many neuroanatomical and neurophysiological characteristics with humans and have long been used as models of aging related cognitive decline [18-23]. Accumulating data indicate aging related neuropathologies like those observed in human AD also occur, suggesting that macaques may be promising models of sporadic AD [24,25]. Cynomolgus macaques (*Macaca fascicularis*), in particular, accumulate age-related cognitive deficits, brain amyloid, and tauopathies [26,27]. Age-related cognitive impairment has been associated with altered cerebrospinal fluid (CSF) biomarkers of AD-like pathology, including decreased beta-amyloid_1-42_ and increased total and phosphorylated tau [28]. Also reminiscent of human health, diabetes mellitus is associated with accelerated age-related brain amyloid accumulation [29]. When fed a Western diet, cynomolgus macaques develop chronic diseases of aging like humans, including coronary and carotid atherosclerosis, obesity, and type 2 diabetes [30]. Macaque society is organized by social status hierarchies, and socially subordinate female macaques develop stress-related pathologies like humans, including coronary and carotid atherosclerosis, obesity, and depression [31], all of which increase AD risk in humans. Thus, these NHPs are appropriate models for investigations of diet and stress effects on the brain.

Here, we report the effects of consumption of a Western-like versus Mediterranean-like diet and psychosocial stress on neuroanatomy in middle-aged, socially housed, dominant and subordinate cynomolgus macaques. We hypothesized that Western diet and the stress of social subordination would result in structural changes in the brain. This long term randomized, controlled preclinical trial had a follow-up period of approximately 31 months. We measured the volumes of several whole brain and individual regions of interest, and cortical thicknesses and volumes in temporoparietal areas relevant to AD neuropathology [32]. We observed widespread neuroanatomical changes in the Western group and psychosocial stress effects on several regions relevant to AD, as well as temporally associated brain changes in all subjects likely due to aging. This work provides important insights into the effects of diet, psychosocial stress, and aging in the primate brain.

## 2. METHODS

### 2.1 Animals and Design

Middle-aged (11-13 years of age) female cynomolgus macaques were obtained (SNBL USA, Alice, Texas) and single-cage quarantined for one month in a room with visual and auditory contact, a routine procedure in animal facilities. Individuals were then moved to small social groups (N=4), housed in 3 × 3 x 3 m^3^ indoor enclosures with exposure to natural light and a 12/12 light-dark cycle (light phase: 0600-1800) and water *ad libitum*. During the period prior to the dietary intervention, animals received monkey chow.

Following a seven-month Baseline Phase, 42 monkeys were assigned to the two experimental diet groups (Mediterranean versus Western). The diet groups were balanced on age (X _MED_=12.1 y; range=11.1-13.6); X _WEST_=12.2 y; range=11.4-13.0), body weight, body mass index, age, basal cortisol concentration, and plasma lipid concentration as previously described [33]. See Supplementary Figure 1 for a schematic of the experimental design.

### 2.2 Ethics Statement

All procedures involving animals complied with the National Institutes of Health Guide for the Care and Use of Laboratory Animals (NIH Publications No. 8023, revised 1978), state, and federal laws, and were approved by the Animal Care and Use Committee of Wake Forest School of Medicine.

### 2.3 Social Status

During the Baseline Phase, animals adapted to their social groups and established stable social status hierarchies. Social status was unknown prior to group formation. Animals that were in the top half of their hierarchy, on average over the course of the experiment, were considered dominant, and all others subordinate. Dominants and subordinates were similar in age (X _SUB_=11.9 y; range=11.1-12.2); X _DOM_=12.3 y; range=11.1-13.6). As previously observed, social status hierarchies were stable across the duration of the experiment, with 92.5% of subjects maintaining their rank through the 31 months [34]. Subordinates were considered stressed because they received more aggression, were groomed less, spent more time alone, and had higher heart rates and cortisol concentrations than dominants (Shively et al., in press).

### 2.4 Experimental Diets

Macronutrient content of the diets is shown in Supplementary Table 1. The experimental diets have been previously described and differed considerably from monkey chow [33]; (Supplementary Note 1). The Western diet was designed to mimic the diet typically consumed by American women (age 40-49), whereas the Mediterranean diet recapitulated key aspects of the human Mediterranean diet [7]. The experimental diets were matched on macronutrients and cholesterol content but differed significantly in fatty acids. Fats and proteins were mostly plant based in the Mediterranean diet [7], whereas fats and proteins were derived mostly from animal sources in the Western diet. This resulted in a high percentage of monounsaturated fats in the Mediterranean diet and saturated fats in the Western diet [7,35]. The Mediterranean diet was higher in complex carbohydrates and fiber, and had a lower n-6:n-3 fatty acid ratio similar to a traditional hunter-gatherer type diet [35], and lower sodium and refined sugars than the Western diet. Key Mediterranean ingredients included English walnut powder and extra-virgin olive oil which were the primary components provided to participants in the PREDIMED study [36]. Changes in body weight and composition resulting from the dietary intervention have been previously reported, and social status was not related to body weight or body composition [33].

**Table 1.** Irrespective of diet or social status, several UNC ROI volumes changed over the course of the experiment. These ROIs included GM, WM, and CSF. Full models were ANCOVAs with repeated measures (2_Mediterranean, Western_ X 2_Dom, Sub_ X 2_Left, Right_ X 2_Base, Exp_). Means have been adjusted for inter-individual variation in intracranial volumes, and the percent change for individual ROIs are shown. Right and left ROIs were analyzed and reported separately if an interaction effect of side was detected. F and p-values for the main effects of time are shown.

|  | UNC ROI | Adj.<br>Base.<br>(mm<sup>3</sup>) | Adj.<br>Exp.<br>(mm<sup>3</sup>) | Percent<br>Change | F | DF1 | DF2 | Effect<br>of<br>Time *p* |
| --- | --- | --- | --- | --- | --- | --- | --- | --- |
| Whole<br>ROIs | Whole<br>Hippocampus | 350.50 | 351.29 | 0.23 | 0.239 | 1 | 112 | 0.626 |
|  | Anterior<br>Hippocampus | 164.89 | 164.65 | -0.15 | 0.034 | 1 | 112 | 0.854 |
|  | Posterior<br>Hippocampus | 185.66 | 186.66 | 0.54 | 0.762 | 1 | 112 | 0.385 |
|  | Temporal Auditory | 2215.63 | 2201.45 | -0.64 | 1.236 | 1 | 112 | 0.269 |
|  | Temporal Visual | 4365.39 | 4306.99 | -1.34 | 11.541 | 1 | 112 | 0.001 |
|  | Prefrontal | 2896.04 | 2846.02 | -1.73 | 32.383 | 1 | 112 | <0.001 |
|  | Frontal | 4080.81 | 4035.76 | -1.10 | 26.174 | 1 | 112 | <0.001 |
|  | Insula | 312.75 | 308.10 | -1.49 | 6.700 | 1 | 112 | 0.011 |
|  | Amygdala | 280.39 | 280.29 | -0.04 | 0.004 | 1 | 112 | 0.947 |
|  | Cingulate | 808.26 | 814.21 | 0.74 | 2.915 | 1 | 112 | 0.091 |
|  | Caudate | 381.15 | 385.62 | 1.17 | 10.034 | 1 | 112 | 0.002 |
|  | Putamen | 479.07 | 484.47 | 1.13 | 6.086 | 1 | 112 | 0.015 |
|  | Parietal | 4061.76 | 4010.69 | -1.26 | 11.811 | 1 | 112 | 0.001 |
|  | Occipital | 5435.93 | 5360.06 | -1.40 | 18.362 | 1 | 112 | <0.001 |
|  | Cerebellum | 3046.96 | 3037.19 | -0.32 | 1.052 | 1 | 112 | 0.307 |
|  | Pons and Medulla | 1117.04 | 1088.98 | -2.51 | 43.277 | 1 | 112 | <0.001 |
|  | Corpus Callosum | 507.11 | 499.26 | -1.55 | 7.071 | 1 | 112 | 0.008 |
| L & R<br>ROIs | Right Temporal<br>Limbic | 1557.63 | 1542.74 | -0.96 | 12.857 | 1 | 37 | 0.001 |
|  | Left Temporal<br>Limbic | 1466.00 | 1468.31 | 0.16 | 0.196 | 1 | 37 | 0.661 |

### 2.5 Neuroimaging

#### 2.5.1 Image Acquisition

Magnetic Resonance Imaging (MRI) scans were performed during the Baseline Phase and after 31 months of dietary intervention. Animals were transferred to the MRI facility, sedated (ketamine HCl, 10-15 mg/kg body weight), and anesthesia was maintained by isoflurane (3% induction, 1.5% maintenance). We obtained T1-weighted anatomic images using a 3D volumetric MPRAGE sequence (TR=2700 msec; TE=3.39 msec; TI=880 msec; Flip Angle=8 degrees; 160 slices, voxel dimension = 0.5 × 0.5 × 0.5 mm^3.^) using a 3T Siemens Skyra scanner (Siemens, Erlangen, Germany) and a circularly polarized, 32-channel head coil with an internal diameter of 18.4 cm (Litzcage, Doty Scientific, SC).

#### 2.5.2 Image Preprocessing

We first implemented bias field correction, denoising, and intensity normalization using the N4BiasFieldCorrection function of Advanced Normalization Tools (ANTs, Penn Image Computing and Science Laboratory, University of Pennsylvania, PA). Next, an initial whole-head, study specific template (SST) was created from the baseline scans for skull stripping using ANTs (antsMultivariateTemplateConstruction.sh) after rigid-body alignment to a UNC Primate Atlas [37]. We registered the UNC Template brain [37] to the SST brain and transformed the UNC parcellation map, brain mask, and segmentation priors to our SST using deformable registration. The full parcellation consisted of separate definitions for the left and right hemisphere for the subcortical, frontal, prefrontal, cingulate, parietal, occipital, auditory, visual and limbic temporal lobes, as well as the brainstem, corpus callosum, and cerebellum. Cortical labels in the parcellation map included gray matter (GM), white matter (WM), and cerebrospinal fluid (CSF), and subcortical structures, including the hippocampus, amygdala, caudate and putamen.

#### 2.5.3 Image Registration and Tissue Segmentation

To estimate longitudinal transformations, we performed within-subject registration of T1-weighted images across the data acquisition time points using ANTs nonlinear registration algorithm (antsRegistrationSyN.sh [38,39]). We then used longitudinal transformations to warp the Baseline brain masks to the Experimental time-point scans. By warping the parcellation map on SST, we generated individual parcellation maps for brain regions of interest (ROIs) for all time points. We performed an automatic brain segmentation into GM, WM, and CSF using probabilistic tissue maps from tissue priors generated in SST space.

### 2.6 Cortical Volume and Thickness in NHP AD Signature ROIs

In humans, an “AD signature” meta ROI, consisting of multiple ROIs for Alzheimer’s disease pathology, provides more reliable diagnostics of disease progression than do separate ROIs [32,40,41]. While volume-and thickness-based meta ROIs have been developed for humans, none exist for NHPs. Thus, we developed similar volume-and a thickness-based AD signature meta ROIs for NHPs using the angular gyrus, the inferior, middle, and superior temporal gyrus, entorhinal cortex, fusiform cortex, supramarginal gyrus, precuneus, and parahippocampus. We determined the thickness and volume of each ROI using the macaque template from the Scalable Brain Atlas [42] by warping the Atlas to subject’s native T1w MR images. We used the “KellyKapowski” tool included with ANTs 2.2.0. which is a wrapper for the DiReCT tool, a registration-based estimate of cortical thickness [43]. The thickness meta ROI was calculated as the volume-weighted average of the thicknesses of the individual ROIs to take into account the overall size of individual regions. The volume meta ROI was calculated as the sum of all the individual ROIs. All ROIs analyzed are listed in Supplementary Table 2.

### 2.7 Statistical Analysis

For each ROI, we conducted two sets of analyses: 1) Repeated measures ANOVAs to examine changes over the course of the experiment and 2) ANCOVAs with the baseline ROI used as a covariate to provide the best estimate of effect sizes. In our analyses of intracranial (ICV) and total brain (TBV) volumes, we used body weight as a covariate to control for inter-individual variation in body size. For the remaining volumes, we included ICV as a covariate to control for inter-individual variation in head size. Analyses of cortical thickness did not include a control variable because thicknesses are independent of head size [32].

All models included a factor for diet (Mediterranean and Western) and status (dominant and subordinate). In repeated measures analyses, we also included a factor for “time” (Baseline and Experimental). For volumes represented by two hemispheric measurements, we included a factor for “hemisphere” (left and right). We analyzed the resulting full models – i.e., all main and interaction effects – and then eliminated the non-significant (p>0.05) four-way, and then three-way interactions. Next, we inspected models for hemispheric interactions. If present, we analyzed the right and left ROIs separately. If absent, we analyzed the ROI as a single structure with repeated measures. This stepwise approach allowed us to focus on the main and interaction effects of diet and status over time.

## 3. RESULTS

A total of 38 animals are included in the analysis (N_WEST_=21, 11 dominant, 10 subordinate; N_MED_=17, 10 dominant, 7 subordinate). Two animals did not tolerate the experimental diets, were fed standard monkey chow and excluded from analyses reported here, and two animals died during the study (Supplementary Note 2).

Whole brain ROIs included segmentations for GM, WM, CSF, total brain (TBV; TBV = GM + WM), and intracranial volumes (ICV; ICV = GM + WM + CSF). The AD signature meta ROIs included only GM tissue, whereas UNC ROIs included GM, WM and CSF. So, while analyses of the AD signature ROIs focused on GM, analyses of the UNC ROIs illustrated the effects in larger brain regions that included all three types of tissue.

### 3.1 Whole Brain Volume ROIs

#### 3.1.2 Diet Effects

Significant diet by time interactions were observed for several whole brain regions. In the Western group, volumes of CSF (diet x time: F(1,35)=8.859; p=0.005; Tukey p=0.001) and WM (diet x time: F(1,35)=7.826; p=0.008; Tukey p= 0.021) decreased over time, whereas TBV (diet x time: F(1,35)=6.679; p=0.014; Tukey p=0.001) and GM (diet x time: F(1,35)=8.739; p=0.006; Tukey p=0.001) increased over time (Figure 1). ICV did not significantly change over time (diet x time: F(1,35)=0.239; p=0.628). ANCOVAs controlling for baseline measures showed that the Western group had larger TBV (F(1,33)=12.515; p=0.001) and GM (F(1,33)=10.830; p=0.002), and smaller CSF (F(1,33)=8.448; p=0.006) and WM (F(1,33)=6.717; p=0.014) volumes. ICV did not differ between diet groups (F(1,33)=3.047; p=0.090).

**Figure 1.**
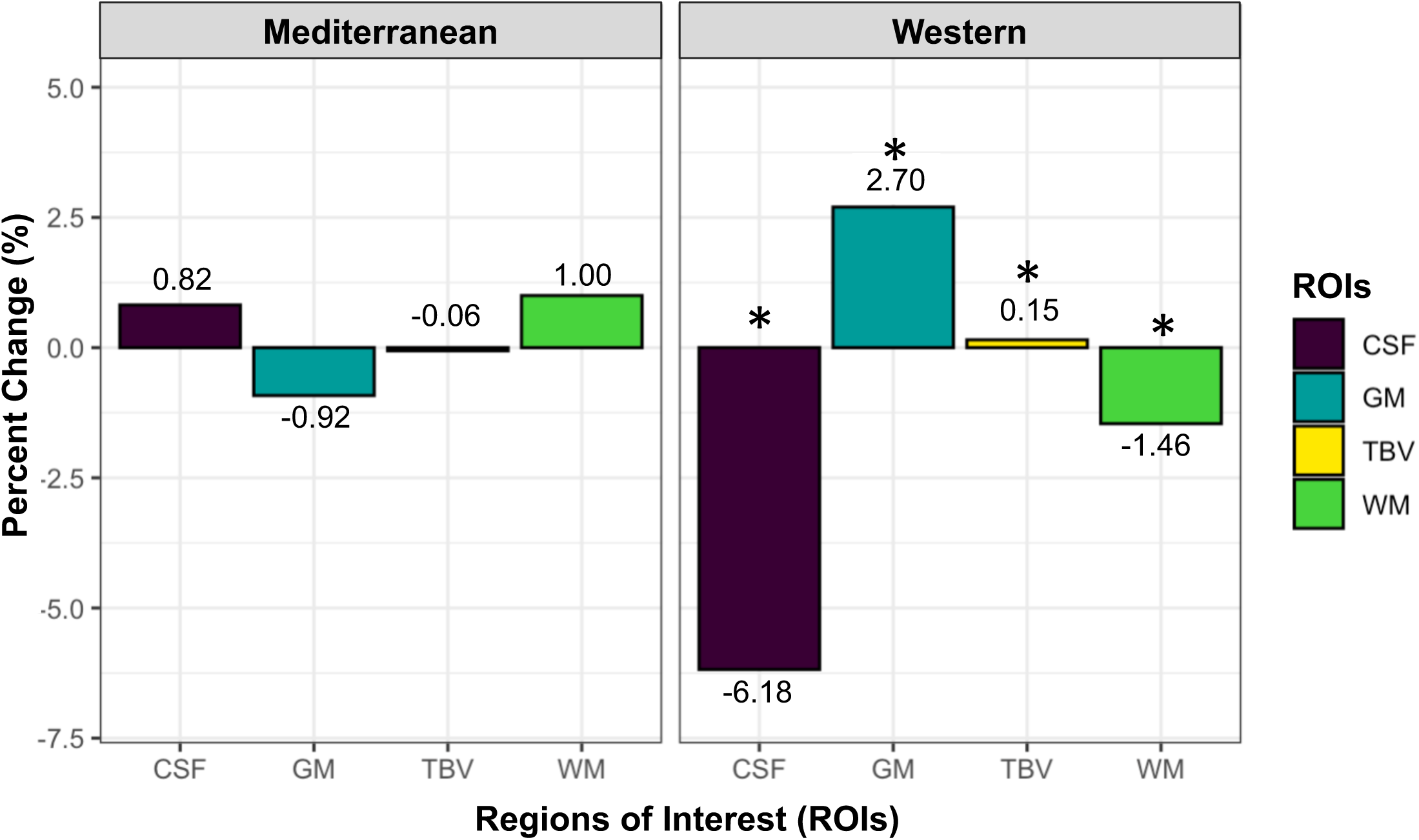
Changes in global brain volumes during experiment. Global brain volumes changed in the Western group, while Mediterranean volumes remained stable over time. In the Western group, total brain (TBV) and gray matter (GM) volumes significantly increased, whereas the volumes of cerebrospinal fluid (CSF) and white matter (WM) significantly decreased. Percent changes in volumes from Baseline to the Experimental time points are shown for each ROI. Significant differences (p≤0.05) indicated by (*).

#### 3.1.3 Status Effects

ICV and TBV differed by social status. After controlling for differences in body size, repeated measures analyses showed that dominants had larger ICV (F(1,34)= 13.805; p=0.001) (Figure 2A) and TBV (F(1,34)=11.497; p=0.002) than subordinates (Figure 2B).

**Figure 2.**
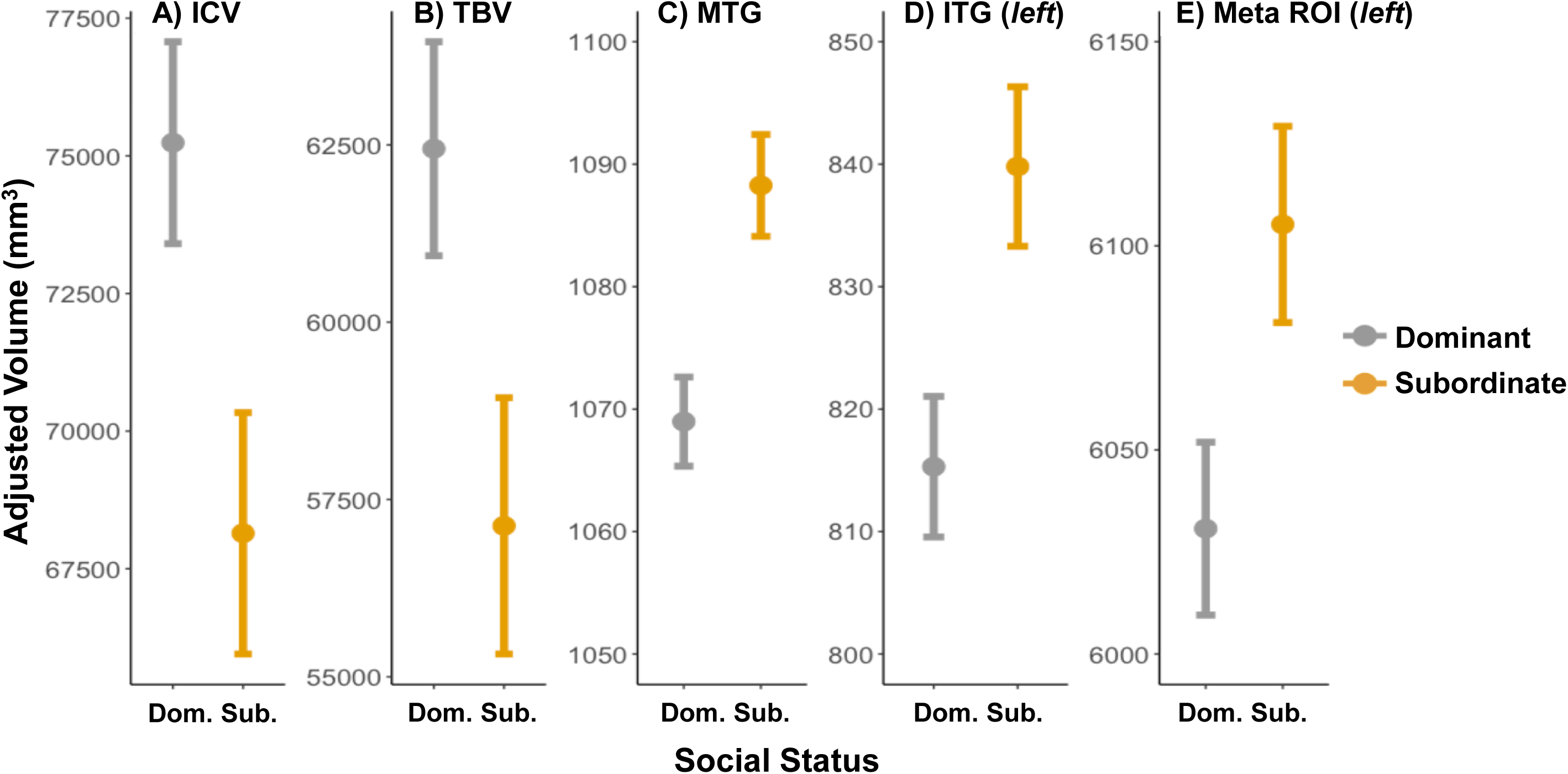
Social status differences in global brain volumes. Socially subordinate monkeys had smaller A) intracranial volumes (ICV) and B) total brain volumes (TBV) than dominants. In contrast, subordinate monkeys had larger volumes than dominants in the following ROIs: C) middle temporal gyri, D) left inferior temporal gyri, and E) left meta-ROI. Adjusted means and standard errors are shown.

#### 3.1.4 Diet by Status Interaction

No significant diet by time interactions were observed for whole brain volume ROIs.

### 3.2 AD Signature ROIs - Cortical Thicknesses

#### 3.2.1 Diet Effects

The entorhinal cortex was the only structure in which the left and right sides differed and for which separate analyses was required. Excluding the right entorhinal cortex (diet x time: F(1,35)=0.012, p=0.914), cortical thicknesses significantly increased in the Western group, whereas cortical thicknesses remained stable over time in the Mediterranean group (Figure 3; Supplementary Table 3).

**Figure 3.**
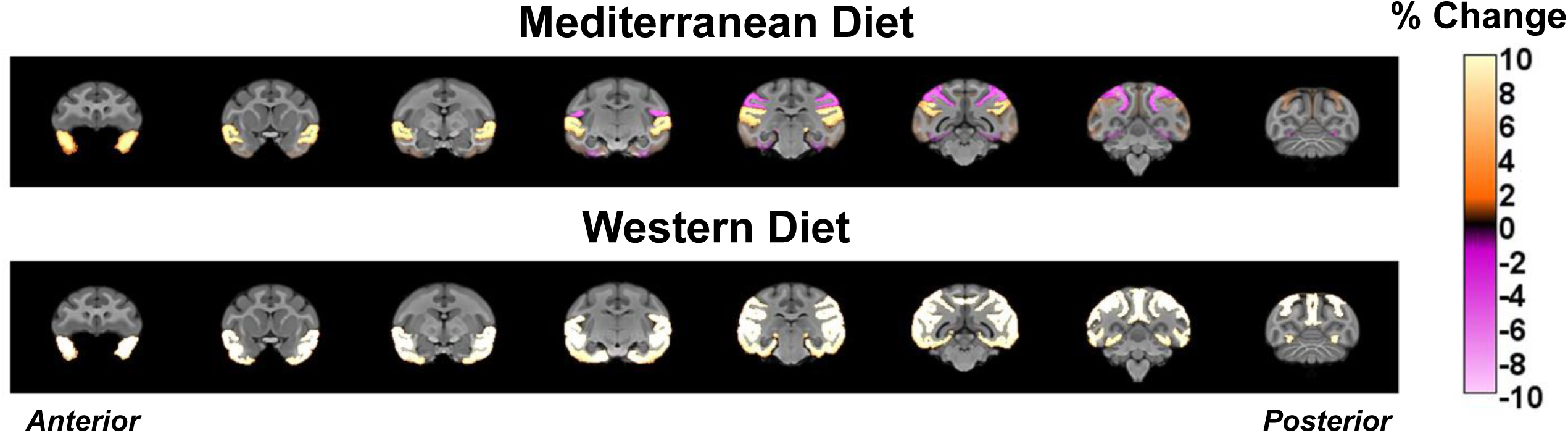
Anatomical map of cortical thickness changes in AD “signature” regions. Cortical thicknesses in regions of interest for AD-like neuropathology increased in animals consuming Western, but not Mediterranean diets. Orange and yellow tones indicate increases in cortical thicknesses, and pink tones indicate decreases in cortical thicknesses. Anterior (left) to posterior (right) coronal sections are shown.

After controlling for inter-individual differences at baseline, ANCOVAs revealed that cortical thicknesses were larger in the Western than the Mediterranean group for most ROIs (Figure 4): angular gyrus (F(1,34)=13.120, p<0.001), inferior temporal gyrus (F(1,34)=8.347, p=0.007), superior temporal gyrus (F(1,34)=18.311, p<0.001), supramarginal gyrus (F(1,34)=16.230, p<0.001), precuneus (F(1,34)=9.702, p=0.003), middle temporal gyrus (F(1,34)=13.211, p=0.001), parahippocampus (F(1,34)=4.4140; p=0.043), and the meta ROI (F(1,34)=14.311, p=0.001). We detected a side by diet interaction effect for the entorhinal cortex. The Western group had larger left entorhinal cortices than the Mediterranean group (F(1,33)=6.316, p=0.017), whereas the right entorhinal cortices did not differ between diet groups (F(1,33)=0.008, p=0.978).

**Figure 4.**
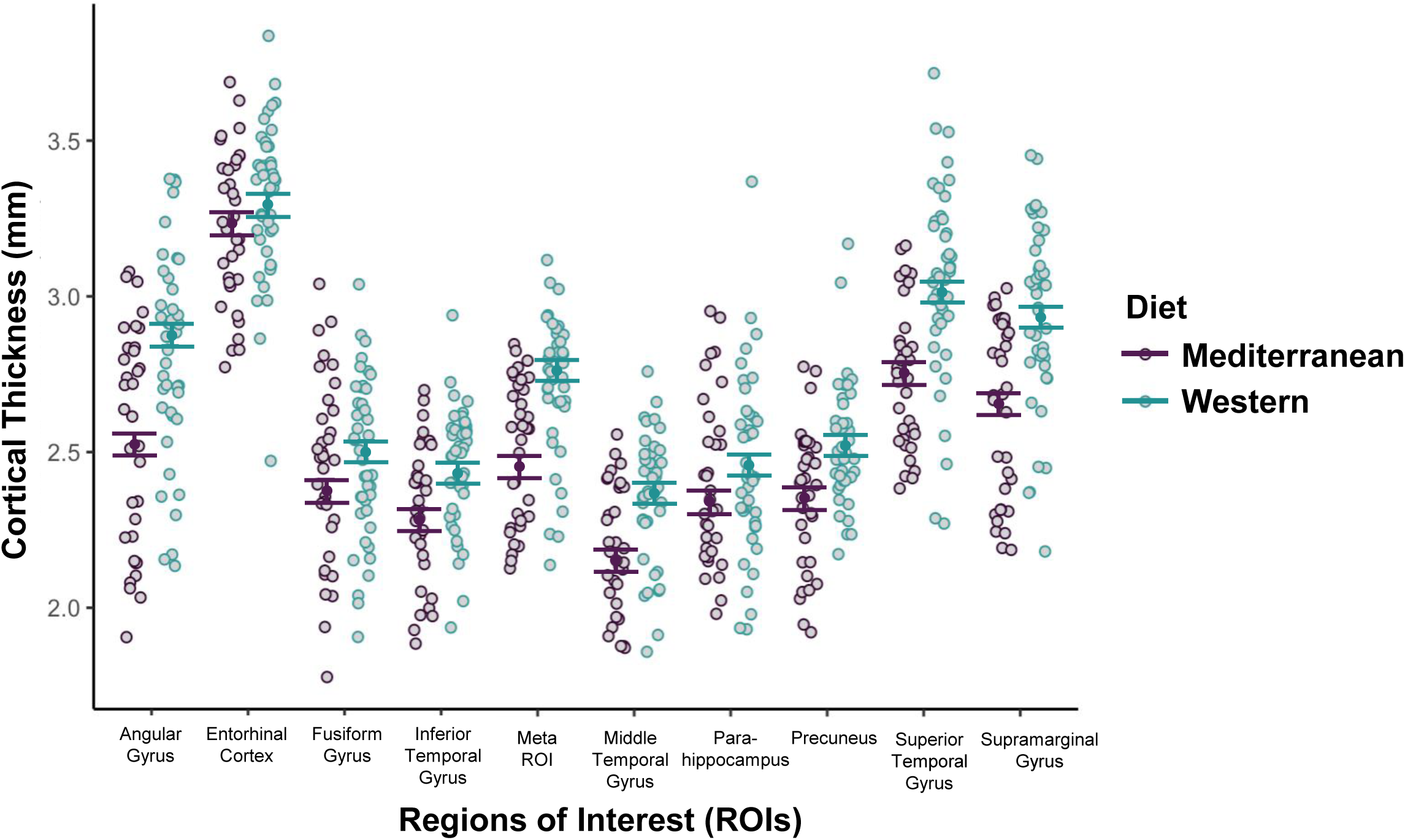
Diet-group differences of cortical thicknesses in AD “signature” regions. Following dietary intervention, Western group cortices were significantly thicker than Mediterranean cortices for every AD-signature region of interest (ROI) except the right entorhinal cortex. Individual values for the ROIs are indicated by the gray points. Adjusted means and standard errors for each diet group are indicated by the solid dots and lines, respectively.

#### 3.2.2 Status Effects

Neither the repeated measures analyses nor the ANCOVAs controlling for baseline revealed any effects of status on cortical thickness.

#### 3.2.3 Diet by Status Interaction

No significant diet by time interactions were observed for the AD-signature cortical thicknesses.

### 3.3 AD Signature ROIs - Cortical Volumes

#### 3.3.1 Diet Effects

Repeated measures analyses of AD-signature cortical volumes revealed significant increases over time in the right inferior temporal gyri (diet x time: F(1,35)=6.606; p=0.015; Tukey=0.006) and right entorhinal cortices (diet x time: F(1,35)=4.582; p=0.039; Tukey=0.365) in the Western group. After controlling for baseline, ANCOVA revealed several main effects of diet. The Western group had larger right inferior temporal gyri (F(1,33)=10.494; p=0.002), right superior temporal gyri (F(1,33)= 5.6090; p=0.024), and right meta ROIs (F(1,33)= 8.179; p=0.007). These ROIs did not differ by diet on the left side of the brain (p’s>0.05; data not shown).

#### 3.3.2 Status Effects

Repeated measures analyses did not detect any significant effects of social status. ANCOVAs indicated that subordinates had larger volumes than dominants in the left and right middle temporal gyri (F(1,35)=6.792; p=0.013), left inferior temporal gyri (F=(1,33)=6.9437; p=0.013), and left meta ROIs (F(1,33)=4.677; p=0.038) (Figure 2C-E).

#### 3.3.3 Diet by Status Interaction

No significant diet by time interactions were observed for the AD-signature cortical volumes.

### 3.4 UNC ROIs

#### 3.4.1 Diet Effects

There were no significant interactions of diet by time for UNC ROIs using repeated measures analyses (p’s>0.05, data not shown). After controlling for baseline, ANCOVA revealed that right caudate volumes were larger in the Western than the Mediterranean group (F(1,33)=4.254; p=0.047) (Figure 5).

**Figure 5.**
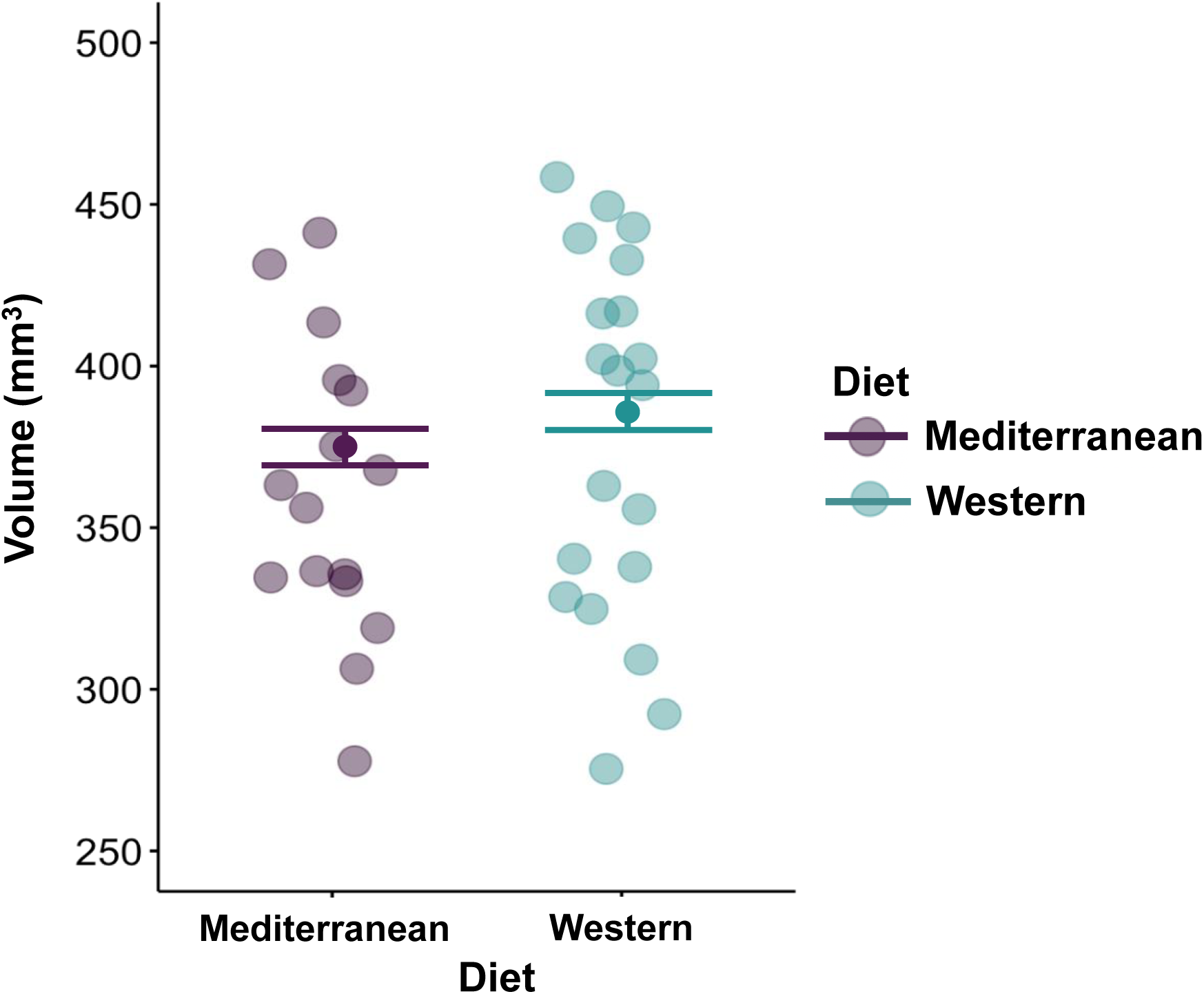
Diet-group differences in caudate volumes. ANCOVA controlling for baseline volumes showed that caudate volumes were smaller in the Mediterranean diet group (377 ± 2.71mm^3^) than in the Western diet group (385 ± 2.40mm^3^) (F(1,33)=4.254; p=0.047). Individual data points are indicated in gray, and adjusted means and standard errors are represented by solid dots and lines, respectively.

#### 3.4.2 Status Effects

Neither the repeated measures analyses nor the ANCOVAs controlling for baseline revealed any main effects of status on any UNC ROI.

#### 3.4.3 Diet by Status Interaction

We detected a diet by status interaction for left frontal volumes (F(1,33)=4.143; p=0.050). Dominants and subordinates did not differ in the Western group (Tukey p=0.662); however, in the Mediterranean group, subordinates had larger frontal volumes than dominants (Tukey p= 0.039) (Figure 6).

**Figure 6.**
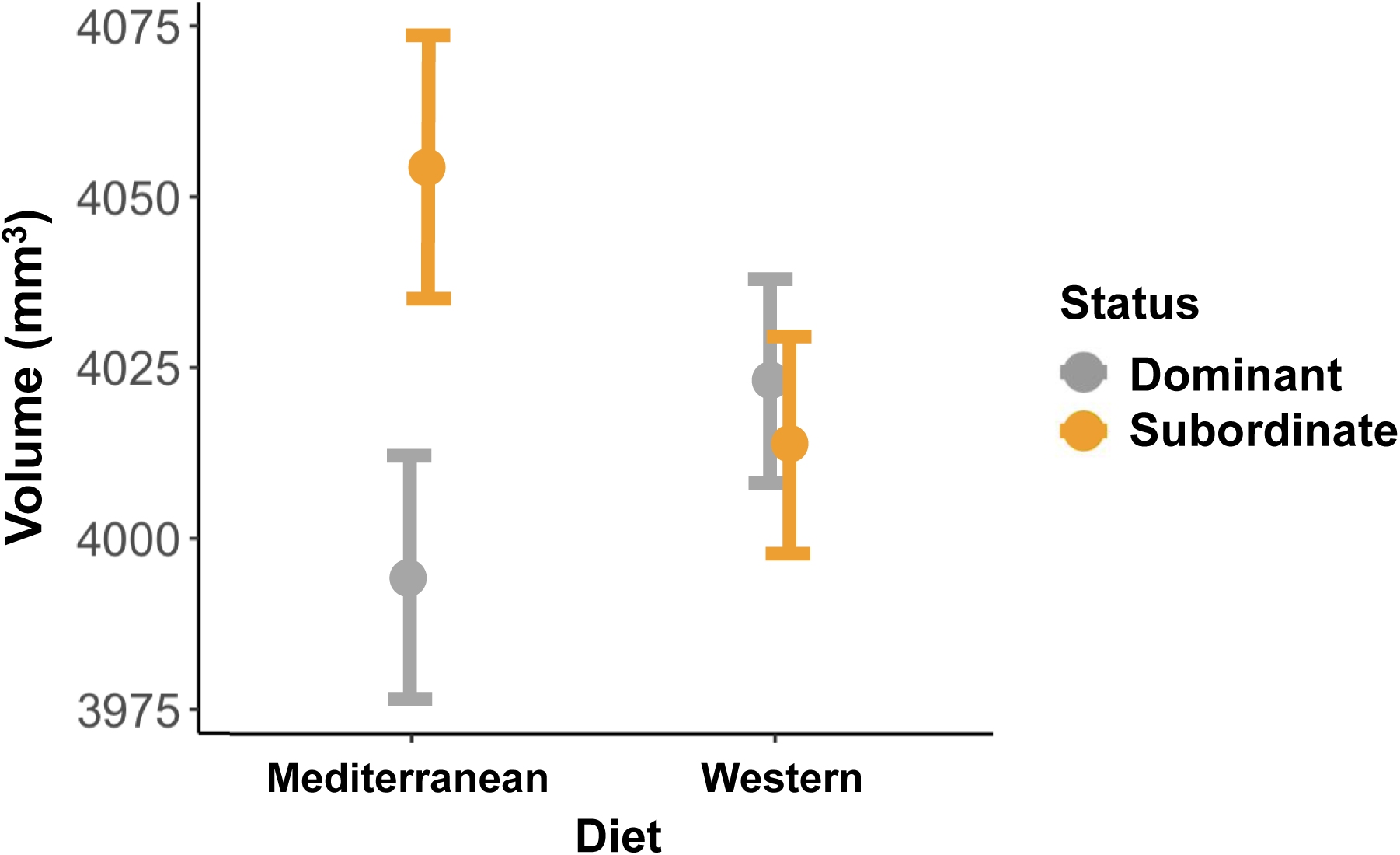
Social status differences in left frontal volumes. In animals consuming Western diets, left frontal volumes were indistinguishable by social status. In contrast, within the Mediterranean diet group, subordinates had significantly larger left frontal volumes than did dominant monkeys (Tukey=0.039). Adjusted means and standard errors after adjusting for baseline volumes are shown.

#### 3.4.4 Time Effects

We detected several main effects of time indicating cohort-wide volumetric changes in ROIs over the 31-month experimental period (Table 1). Most ROIs decreased. However, the caudate (F(1,112)= 10.034; p=0.002) and putamen (F(1,112)=6.086; p=0.015) volumes increased.

## 4. DISCUSSION

The goal of this longitudinal, randomized, dietary intervention was to identify mid-life changes in brain structure due to either stress or diet that are associated with AD risk. Female cynomolgus macaques have an average lifespan of 25 years [44], thus our 11-13 year old cohort were similar in biological age to middle-aged premenopausal women. Middle-age is of particular interest because mounting evidence suggests that structural brain changes in those later diagnosed with AD may begin decades before onset of symptoms [45]. Our findings add to a growing body of evidence that diet and psychosocial stress have profound effects on the brain. Western diet resulted in greater cortical thicknesses, total brain volumes and gray matter, and diminished CSF and white matter volumes. Socially stressed subordinates had smaller whole brain volumes but larger ROIs relevant to AD than dominants. These results provide a foundation upon which future studies can help determine how diet and psychosocial stress influence risk for neurodegenerative disorders such as AD.

### 4.1 Diet Effects

In support of our hypothesis that Western diets promote neuroanatomical changes, the Western group exhibited increased GM volume as well as cortical thicknesses in temporoparietal regions relevant to AD neuropathology. This may see counterintuitive as large GM and cortical volumes are often assumed to indicate a healthy brain. Our cohort consisted of 11 to 13-year-old monkeys, roughly comparable to 35 to 50-year-old humans, which may be too young for development of AD-like neuropathology. However, mounting evidence suggests that early, preclinical increases in GM may portend AD later in life [46-48]. The biological mechanisms underlying increased GM volume and cortical thickening require further study; however, they may reflect neuroinflammatory responses, as increases in GM volumes [49] and cortical thickening [50] have been associated with neuroinflammation in recent clinical studies. Western diets are pro-inflammatory [11,51], and we previously reported that circulating monocytes isolated from the Western group had pro-inflammatory gene expression profiles relative to those from the Mediterranean group [52]. We also reported that the Western group had lower rates of affiliation and social integration and higher rates of anxiety and social isolation than the Mediterranean group, suggesting that diet impacted the brain [52]. Importantly, social isolation and anxiety are risk factors for dementia [53]. Considering these observations, it is possible that the Western diet induced neuroinflammation that promoted macrostructural changes, and that the enlarged cortical and other GM structures observed here reflect an early transition from normal to pathological aging. Chronic inflammation is increasingly recognized as a key mechanism in AD [54]. These observations underscore the need for longitudinal MRI studies in humans that span the range of normal to MCI to AD.

WM volumes decreased in the Western group but remained stable in the Mediterranean group. Changes in WM volumes are some of the earliest biomarkers of AD neuropathology [5,55]. Pathways associated with oxidative damage to oligodendrocytes may play a role in the associations between diet, WM aberrations, and AD [55]. Over time, damage to oligodendrocytes may culminate in the pathogenesis of WM abnormalities and subsequent neurodegenerative disease. The lack of changes in WM in the Mediterranean group may reflect the protective actions of antioxidants in the diet. Recent research in NHPs has shown that consumption of the antioxidant curcumin improves WM integrity [56]. Mediterranean diets are high in polyphenols, compounds that act as both antioxidants and anti-inflammatory agents [10]. The PREDIMED study demonstrated associations of Mediterranean diet with increases in the activity of endogenous antioxidants [57]. Thus, antioxidant properties of the Mediterranean diet may protect WM.

The Western group had larger caudates than the Mediterranean group. In humans, the caudate is part of the salience network, a circuit involved in the regulation of consumptive behavior [58]. We previously reported that the Western group consumed more calories and accumulated higher body fat percentages than the Mediterranean group [30]. These observations suggest that Western diets may alter neural networks in ways that promote over-eating and obesity, an important risk factor for AD [1].

### 4.2 Psychosocial Stress Effects

Stressed subordinates had smaller overall brain volumes and larger volumes of temporoparietal ROIs relevant to AD neuropathology. These results add to a body of literature showing that chronic exposure to psychosocial stress is associated with structural changes in the brain [59]. While this is the first report of social status differences in whole brain volumes in NHPs, much work has reported associations between brain volumes and stress. Subordinates also had volumetric increases in the middle and inferior temporal gyri, and the meta-ROI. Longitudinal studies are needed to determine whether the volumetric differences observed here precede functional impairments. Several plausible mechanisms may underlie these stress-associated volumetric changes, including neuroinflammation [60], neuronal remodeling [61], or neuronal apoptosis [62]. Further studies are needed to elucidate pathways mediating the relationships between psychosocial stress and brain volumes.

### 4.3 Diet –Status Interaction Effects

Left frontal volumes were smaller in dominants than subordinates in the Mediterranean group, whereas they were not different in the Western group, with volumes falling about midway between the Mediterranean dominants and subordinates. It may be that Western diet diminishes differences between dominants and subordinates. Additional study is needed to replicate and explore this observation.

### 4.4 Aging Effects

In addition to the effects of diet and psychosocial stress, several ROI volumes decreased over time. On average, about 2.7 years passed between the MRI scans, a timespan approximating an 8-10 year follow up in humans [44]. These trends may well reflect changes associated with the process of aging. Similar changes in brain volumes have been demonstrated in longitudinal clinical surveys [63]. Changes in autonomic nervous system and hypothalamic-pituitary-adrenal (HPA) function also reflected the aging process in this cohort (Shively et al., in press). These cohort-wide changes are notable because they occurred at ages that roughly translate to 40-49 year old women – a period of time that has been hypothesized to represent a critical period for the development of AD-related neuropathologies [5].

### 4.5 Strengths, Challenges, & Limitations

Strengths of our study include the longitudinal design, careful control of the dietary manipulation, and a focus on a limited number of regions of interest. Notwithstanding these strengths, there were limitations to this study. The Mediterranean diet was developed specifically for this study; thus, this diet was a new formulation. Females were the focus of study because the prevalence of AD is higher in women than in men [17], there are sex differences in etiology [72], and there are sex differences in health responses to social stressors. Thus, a direct comparisons using the same stressor was not scientifically defensible [64]. Repeated longitudinal MRI scans may have more specifically pinpointed the timing of the observed effects but were beyond the scope of this project. Cognitive assessments would provide an opportunity to determine structure-function relationships but were not included as they require food restriction, food reward, and social separation for testing, all of which were incompatible with our study goals and design. Future studies are planned to replicate these observations in males, conduct scans more frequently, and assess cognitive function.

### 4.6 Conclusions

These results suggest that persistent psychosocial stress and Western diet consumption result in macrostructural changes in the brain. The associations between Western diet and brain changes may be due, in part, to changes in proinflammatory pathways that induce structural changes in the brain. In contrast, the anti-inflammatory and antioxidant properties of Mediterranean diets may underlie their protective effects. However, these hypotheses need to be confirmed with further neurobiological phenotyping, which is underway. Future studies using innovative molecular approaches, alternative imaging modalities, and novel biomarkers are necessary to characterize the temporal emergence of neuropathological changes. Such work will help to identify resilient versus at-risk phenotypes during middle age, so that intervention can occur before the development of mild cognitive impairment, AD, and dementia.

## ACKNOWLEDGMENTS

This work was supported by the NIH (R01-HL087103 (CAS), RF1-AG058829 (CAS & SC), R01-HL122393 (TCR), and U24-DK097748 (TCR), an Intramural Grant from the Department of Pathology, the Wake Forest Alzheimer’s Disease Research Center (P30-AG049638) and the Wake Forest Claude D. Pepper Older Americans For Independence Center (P30 AG21332). The authors also would like to acknowledge the contributions of the Wake Forest Clinical and Translational Science Institute.

## DECLARATIONS OF INTEREST

none

## SUPPLEMENTARY INFORMATION

**Supplementary Note 1:** A standard monkey lab chow diet group was not included because it is unlike the diet that monkeys consume in the wild [65], there is no human diet analog, it is very low in fat (13%), and the predominant protein source is soy which is rich in isoflavones (diet 5037/5038, LabDiet, St. Louis, MO; see Supplementary Table 1). Isoflavones are known selective estrogen receptor modulators, with independent effects on the brain that have been extensively described by our lab and others in rodent and in primate models (e.g., [66,67]).

**Supplementary Note 2:** While most of the monkeys did well on the diets, two animals died. They were both in the Mediterranean group and both the most socially subordinate in their groups. Both had diarrhea, the cause of which remained undiagnosed in spite of extensive testing. Compared to Baseline, monkeys in the Mediterranean group did not gain weight but those on the Western diet did [33]. Since they were thinner, they had less of a cushion to weather a bout of diarrhea. Social subordination stress may also have been a contributing factor.

**Supplementary Figure 1.**
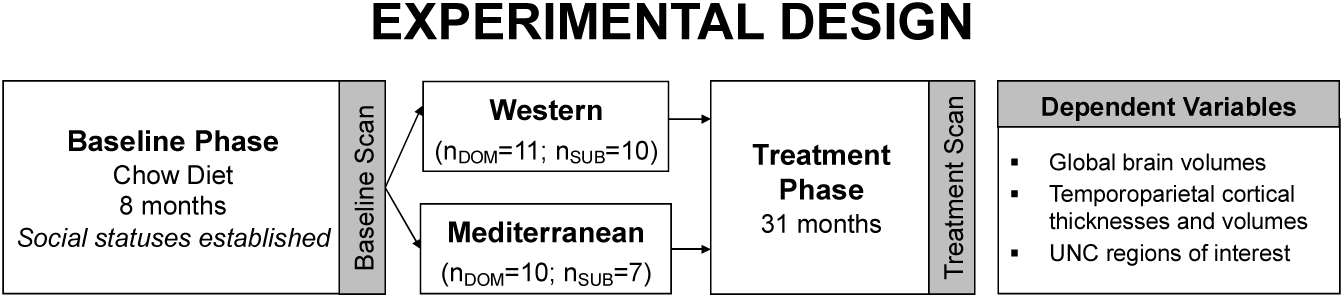
Schematic representation of the experimental design, including the timing of the MR scans, sample sizes, and dependent variables measured.

**Supplementary Table 1.**
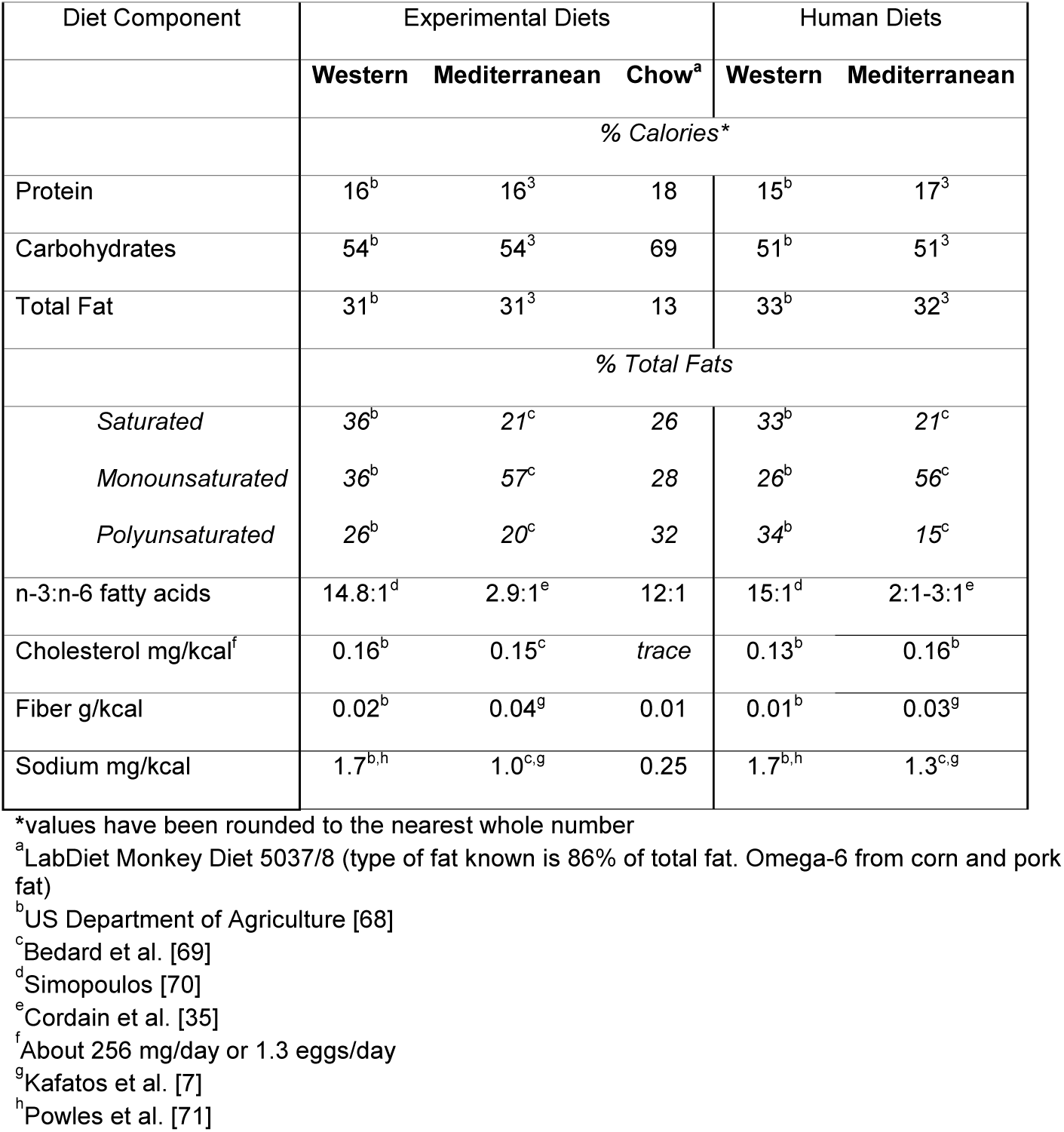
Macronutrients in the Western, Mediterranean, and monkey chow experimental diets, compared to corresponding human diets (from [33]).

**Supplementary Table 2.** Regions of interest (ROIs).

| AD-Signature ROIs<sup>*a</sup> | Whole Brain Volumes<sup>b</sup> | UNC ROIs<sup>b</sup> |
| --- | --- | --- |
| Angular gyrus | Intracranial volume (ICV) | Anterior hippocampus |
| Inferior temporal gyrus | Total brain volume (TBV) | Posterior hippocampus |
| Middle temporal gyrus | Gray matter (GM) | Whole hippocampus |
| Inferior temporal gyrus | White matter (WM) | Temporal auditory |
| Precuneus | Cerebrospinal fluid (CSF) | Temporal visual |
| Fusiform gyrus |  | Temporal limbic |
| Supramarginal gyrus |  | Occipital |
| Entorhinal cortex |  | Frontal |
| Parahippocampal gyrus |  | Insula |
| Meta ROI |  | Cingulate |
|  |  | Parietal |
|  |  | Prefrontal |
|  |  | Corpus Callosum |
|  |  | Pons & Medulla |
|  |  | Amygdala |
|  |  | Caudate |
|  |  | Putamen |

*\*Cortical thicknesses (mm) and cortical volumes (mm<sup>3</sup>) generated for these regions of interest*
a. Neuromaps Scalable Atlas [42]
b. UNC Primate Atlas [37]

**Supplementary Table 3.**
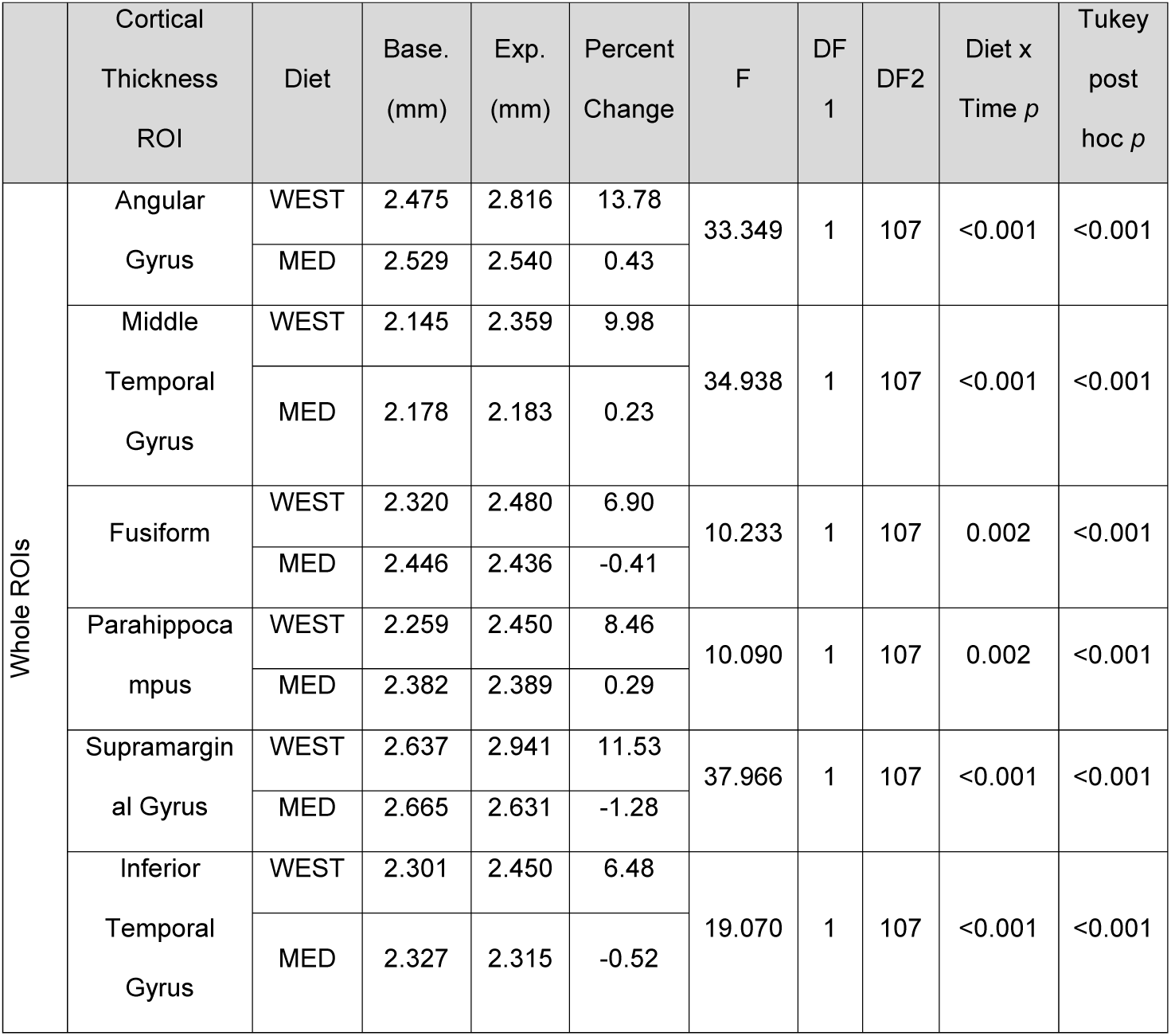
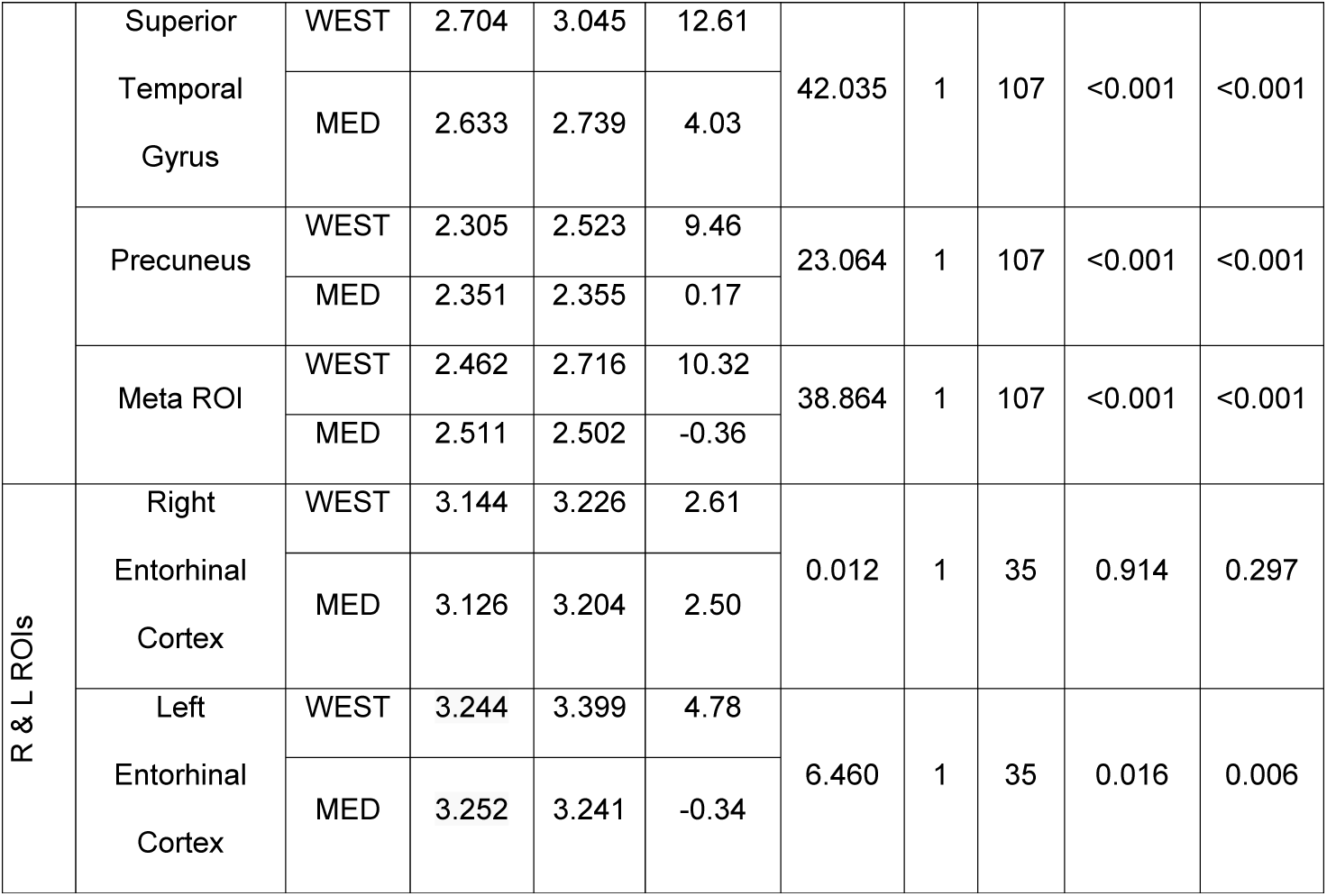
Diet-induced changes in cortical thickness regions of interest (ROI) for AD-like neuropathology. Cortical thicknesses in animals consuming Western diets increased over time, whereas the cortical thicknesses of Mediterranean animals were generally stable over time. Full models were ANOVAs with repeated measures (2_Mediterranean, Western_ X 2_Dom, Sub_ X 2_Left, Right_ X 2_Base, Exp_). P-values corresponding to diet by time interactions are shown. Post hoc *p* values were adjusted using the Tukey method for comparing a family of 4 estimates and indicate differences across time points within the Western diet group.

|  | Cortical Thickness ROI | Diet | Base. (mm) | Exp. (mm) | Percent Change | F | DF 1 | DF2 | Diet x Time *p* | Tukey post hoc *p* |
| --- | --- | --- | --- | --- | --- | --- | --- | --- | --- | --- |
| Whole ROIs | Angular Gyrus | WEST | 2.475 | 2.816 | 13.78 | 33.349 | 1 | 107 | <0.001 | <0.001 |
|  |  | MED | 2.529 | 2.540 | 0.43 |  |  |  |  |  |
|  | Middle Temporal Gyrus | WEST | 2.145 | 2.359 | 9.98 | 34.938 | 1 | 107 | <0.001 | <0.001 |
|  |  | MED | 2.178 | 2.183 | 0.23 |  |  |  |  |  |
|  | Fusiform | WEST | 2.320 | 2.480 | 6.90 | 10.233 | 1 | 107 | 0.002 | <0.001 |
|  |  | MED | 2.446 | 2.436 | -0.41 |  |  |  |  |  |
|  | Parahippocampus | WEST | 2.259 | 2.450 | 8.46 | 10.090 | 1 | 107 | 0.002 | <0.001 |
|  |  | MED | 2.382 | 2.389 | 0.29 |  |  |  |  |  |
|  | Supramarginal Gyrus | WEST | 2.637 | 2.941 | 11.53 | 37.966 | 1 | 107 | <0.001 | <0.001 |
|  |  | MED | 2.665 | 2.631 | -1.28 |  |  |  |  |  |
|  | Inferior Temporal Gyrus | WEST | 2.301 | 2.450 | 6.48 | 19.070 | 1 | 107 | <0.001 | <0.001 |
|  |  | MED | 2.327 | 2.315 | -0.52 |  |  |  |  |  |
|  | Superior<br>Temporal<br>Gyrus | WEST | 2.704 | 3.045 | 12.61 | 42.035 | 1 | 107 | <0.001 | <0.001 |
|  |  | MED | 2.633 | 2.739 | 4.03 |  |  |  |  |  |
|  | Precuneus | WEST | 2.305 | 2.523 | 9.46 | 23.064 | 1 | 107 | <0.001 | <0.001 |
|  |  | MED | 2.351 | 2.355 | 0.17 |  |  |  |  |  |
|  | Meta ROI | WEST | 2.462 | 2.716 | 10.32 | 38.864 | 1 | 107 | <0.001 | <0.001 |
|  |  | MED | 2.511 | 2.502 | -0.36 |  |  |  |  |  |
| R & L ROIs | Right<br>Entorhinal<br>Cortex | WEST | 3.144 | 3.226 | 2.61 | 0.012 | 1 | 35 | 0.914 | 0.297 |
|  |  | MED | 3.126 | 3.204 | 2.50 |  |  |  |  |  |
|  | Left<br>Entorhinal<br>Cortex | WEST | 3.244 | 3.399 | 4.78 | 6.460 | 1 | 35 | 0.016 | 0.006 |
|  |  | MED | 3.252 | 3.241 | -0.34 |  |  |  |  |  |

## Notes

### Competing Interest Statement

The authors have declared no competing interest.

